# Continuous lesion images drive more accurate predictions of outcomes after stroke than binary lesion images

**DOI:** 10.1101/2024.10.04.616726

**Authors:** Thomas M.H. Hope, Douglas Neville, Mohamed L. Seghier, Cathy J. Price

**Affiliations:** Department of Imaging Neuroscience, Institute of Neurology, University College London, 12 Queen Square, London, WC1N 3AR, United Kingdom; Department of Psychological and Social Sciences, John Cabot University, Via della Lungara 233, Rome, 00165, Italy; Department of Biomedical Engineering & Biotechnology, Khalifa University of Science and Technology, Abu Dhabi, UAE

**Keywords:** stroke, language, cognition, machine learning, lesions

## Abstract

Current medicine cannot confidently predict who will recover from post-stroke impairments. Researchers have sought to bridge this gap by treating the post-stroke prognostic problem as a machine learning problem. Consistent with the observation that these impairments are caused by the brain damage that stroke survivors suffer, information concerning where and how much lesion damage they have suffered conveys useful prognostic information for these models. Much recent research has considered how best to encode this lesion information, to maximise its prognostic value. Here, we consider an encoding that, while not novel, has never before been formally examined in this context: ‘continuous lesion images’, which encode continuous evidence for the presence of a lesion, both within and beyond what might otherwise be considered the boundary of a binary lesion image. Current state of the art models employ information derived from binary lesion images. Here, we show that those models are significantly improved (i.e., with smaller Mean Squared Error between predicted and empirical outcome scores) when using continuous lesion images to predict a wide range of cognitive and language outcomes from a very large sample of stroke patients. We use further model comparisons to locate the predictive advantage to the provision of continuous lesion evidence beyond the boundary of binary lesion images. The continuous lesion images thus provide a straightforward way to incorporate details of both lesioned and non-lesioned tissue when predicting outcomes after stroke.

## Introduction

Stroke is common, and those who survive the initial insult often suffer from cognitive impairments. Language impairments (aphasia) occur in around 1/3 of these patients, and can be especially distressing [1]; these patients want to know whether and when they might recover. Recently, many researchers have sought to answer these questions by learning associations between prognostic factors and outcomes, for patients whose outcomes are known, and generalising those trends to predict outcomes for new patients (e.g., [2-5]). Information extracted from brain imaging, concerning where and how much brain lesion damage these patients have suffered has proved useful for this work. However, it remains unclear how best to encode this information to maximise its prognostic value [2-5]. Here, we test how prediction models benefit from using brain images that convey a continuous measure of whole brain structural integrity, rather than a binary encoding that simply distinguished ‘lesioned’ from ‘preserved’ tissue.

The use of continuous images for lesion analysis was first proposed by Stamatakis and Tyler [11] who conducted a group comparison of unsegmented whole brain T1 images from (A) patients with lesions and (B) neurotypical controls. The disadvantages of using unsegmented images are that (i) brain differences might pertain to features that are not related to the brain lesion and (ii) the comparison of lesion sites in different groups of patients is only possible in reference to groups of neurotypical controls. Our preferred method, described in the next paragraph, is to create a continuous lesion image for each patient which represents how much brain structure in each patient differs from that of a control group; without imposing a binary lesion vs non-lesion threshold. We have previously discussed the advantage of using continuous versus binary lesion images in Gajardo Vidal et al., [12]. Others have used continuous lesion images in stroke outcome prediction models [13, 14], but as far as we know, no one has yet attempted to quantify the predictive benefits that can accrue from them, relative to variables derived from binary lesion images. This is our focus in what follows.

Continuous lesion images are best understood by comparison to more familiar binary lesion images, which simply discriminate voxels that are thought to be occupied by a lesion (‘1’) from voxels that are thought to be preserved (‘0’). This binary distinction has typically been made “by an expert eye”, who manually draws lesion boundaries directly onto structural brain images. Alternatively, binary images might be created by more automated procedures [15, 16]. By contrast, continuous lesion images represent the evidence that a lesion occupies each voxel: e.g., a continuous variable in the range 0-1, where ‘1’ indicates maximal evidence for the presence of a lesion, and ‘0’ indicates maximal evidence against the presence of a lesion.

This continuous information is an interim output of our standard MRI pre-processing pipeline for the Predicting Language Outcomes and Recovery After Stroke (PLORAS) data, which derives these numbers by comparing the patient’s scan to a distribution of scans acquired from (age-matched) neurologically normal controls [17]. Binary lesion images can be derived from continuous lesion images via thresholding – as we have done in much of our own past work [2, 3, 18, 19]. This aligns our work with others using binary image inputs derived from manual segmentations. But continuous lesion images encode more information than binary lesion images, including continuous variation in lesion evidence beyond what might otherwise be considered the ‘binary lesion boundary’ and beyond what can be determined by eye. Anecdotally, we have long wondered if the loss of this continuous information, in the conversion from continuous to binary lesion images, might significantly reduce our models’ predictive performance. Here, we answer this question empirically, and find that the loss is significant.

## Methods

### Data

Our patient data are drawn from the Predicting Language Outcomes and Recovery After Stroke (PLORAS) dataset [20], which associates more than 1,300 stroke survivors with: (a) clinical and demographic data; (b) high resolution, T1-weighted structural MRI; and (c) scores from the Comprehensive Aphasia Test (CAT) [21]: a standardised battery of behavioural tasks, designed to assess the severity of participants’ language and cognitive impairments. Data were acquired primarily but not exclusively in the chronic phase, from 2 months to >10 years post-stroke. Patients were recruited into the study over more than ten years. We have described this dataset in detail elsewhere [20]; here, we repeat only those elements that are salient to the current analysis. Our sample excludes patients from this dataset when: (a) English was not their native language; or (b) our MRI pre-processing pipeline (described later on in this section) identified no (binary) lesion from their scan. Patients were included in the PLORAS study if they have suffered a stroke, and could be scanned safely with MRI at our scanners in central London. Patients were excluded from the PLORAS study if they suffered any significant neurological disorder beyond the stroke itself (such as Parkinson’s or Alzheimer’s disease). The data are curated by the PLORAS patient team, including those who acquired the data and have access to identifiable information on the participants. For the purposes of this study, the authors accessed an anonymized version of the dataset on the 14^th^ of December 2023.

The CAT defines 34 task scores, including 29 that refer to language skills, and 5 that refer to ostensibly non-linguistic, general cognitive skills [21]. The latter include a line bisection task (used to assess hemispatial neglect), a semantic memory task (matching pictures of objects / animals on the basis of semantic associations), a recognition memory task (testing memory for items presented in the semantic memory task), a gestural task requiring participants to recognise common objects by miming their use, and a mental arithmetic task. In what follows, we consider attempts to predict all of these 5 “cognitive” task scores, along with 8 “summary” scores that each capture overall performance on a group of similar tasks (as described in the CAT). Specifically, these scores refer to patients’ skills in: (a) fluency tasks; (b) the comprehension of auditory language; (c) the comprehension of written language; (d) the repetition of heard language; (e) naming; (f) the verbal description of visually presented scenes; (g) reading; and (h) writing. Taken together, the 8 language summary scores provide a reasonable profile of participants’ language skills. Therefore, 13 outcome variables (5 non-linguistic and 8 language summary scores) are included in the analyses.

Our T1-weighted MRI scans were acquired using a variety of Siemens scanners, typically but not exclusively on the same day as the behavioural assessment, or within a few days of it. All scans were processed using the Automatic Lesion Identification (ALI) toolbox [17], which is an elaboration of the popular Unified Segmentation algorithm [22], adapted for use in the damaged brain. The ALI toolbox derives continuous lesion evidence at the voxel level by comparing each participant’s scan to a distribution of reference scans, acquired on the same scanners from neurologically normal controls. The result is a whole-brain continuous lesion image, in standard Montreal Neurological Institute space. Binary lesion images are thresholded continuous lesion images (threshold = 0.3).

### Analysis

This is a model comparison study, aiming to measure whether a new type of model (driven by continuous lesions) is better than a baseline model (driven by binary lesions) when predicting language outcomes after stroke. Our baseline is the model that we have found to be most effective in prior studies that considered subsets of the same patient data [2, 3, 18]: these models encode lesion images as ‘lesion load’ variables. Specifically, we re-encode both the binary and the continuous lesion images as lesion load variables: i.e., representing the mean lesion signal within a series of 398 anatomical region masks, extracted from publicly available atlases [23-26]. To these lesion load variables, we append further variables representing: (a) the time post-stroke at which participants were scanned; (b) their age; (c) their pre-stroke handedness; (d) their sex assigned at birth; and their lesion volume in each of the (e) left and (f) right hemispheres assessed from the binary lesion images.

Our analyses then train and test machine learning models driven by these predictors (i.e., one analysis with lesion load variables derived from binary lesion images, and another with predictors derived from continuous lesion images), and compare the resulting Root Mean Squared Errors (RMSE). When other details of the analysis – the model type, and train/test folds for cross-validation – were held constant, we expected that models driven by the continuous lesion images would yield smaller magnitude RMSEs.

We also created two further sets of lesion load variables, from continuous lesion images, adjusted by removing continuous information either: (a) inside; or (b) outside the binary lesion boundary. Images including continuous lesion information only beyond the binary lesion boundary are referred to as ‘continuous-out’; in this case, all voxel values inside the binary lesion boundary are set to ‘1’. Images including that information only within the binary lesion boundary are referred to as ‘continuous-in’; here, all voxels outside the binary lesion boundary are set to ‘0’. The goal here was to try to locate the source of any predictive advantage conveyed by the continuous lesion images, with the expectation that both types of information will have an advantage [27-29] All prediction comparisons were made as described above for binary versus continuous lesion images.

Our model comparison procedure follows the recommendations made by Dietterich [30]: 5 repetitions of a 2-fold (split-half) cross-validation, with consistent folds across models, followed by a pair-wise comparison of the resultant prediction error measures, which we implement with Wilcoxon signed ranks tests.

To account for potential inducer-dependence in our results [3], we ran the whole analysis twice, using each of two different machine learning models. The first is a linear support vector machine, which has been used to good effect by others in the recent past when making analogous predictions [5]. And the second is a bagged random forest, which has proved effective when making these predictions on subsets of the current sample in the past [3]. Both models’ hyperparameters are left at their default values, as specified by Matlab 2023a.

## Results

The random forest model yielded significantly better predictive performance (smaller magnitude prediction errors, RMSEs) than the linear support vector machine, for every outcome variable (all p < 0.001). Table 1 (for the linear support vector machines) and Table 2 (for the random forests) compare the prediction errors that result when the same models are used to predict the same outcome variables, using either: (a) binary lesion images; (b) continuous lesion images; (c) continuous-in images; or (d) continuous-out images. When using a support vector machine, models driven by continuous lesion images always significantly outperformed models driven by binary lesion images: i.e., there was a significant ‘continuous-binary advantage’ for all 13 outcome variables. When using a random forest, there was a significant continuous-binary advantage for 11/13 outcome variables, and a numerical (non-significant) advantage for the other two models (measuring participants’ semantic memory and repetition skills).

**Table 1:**
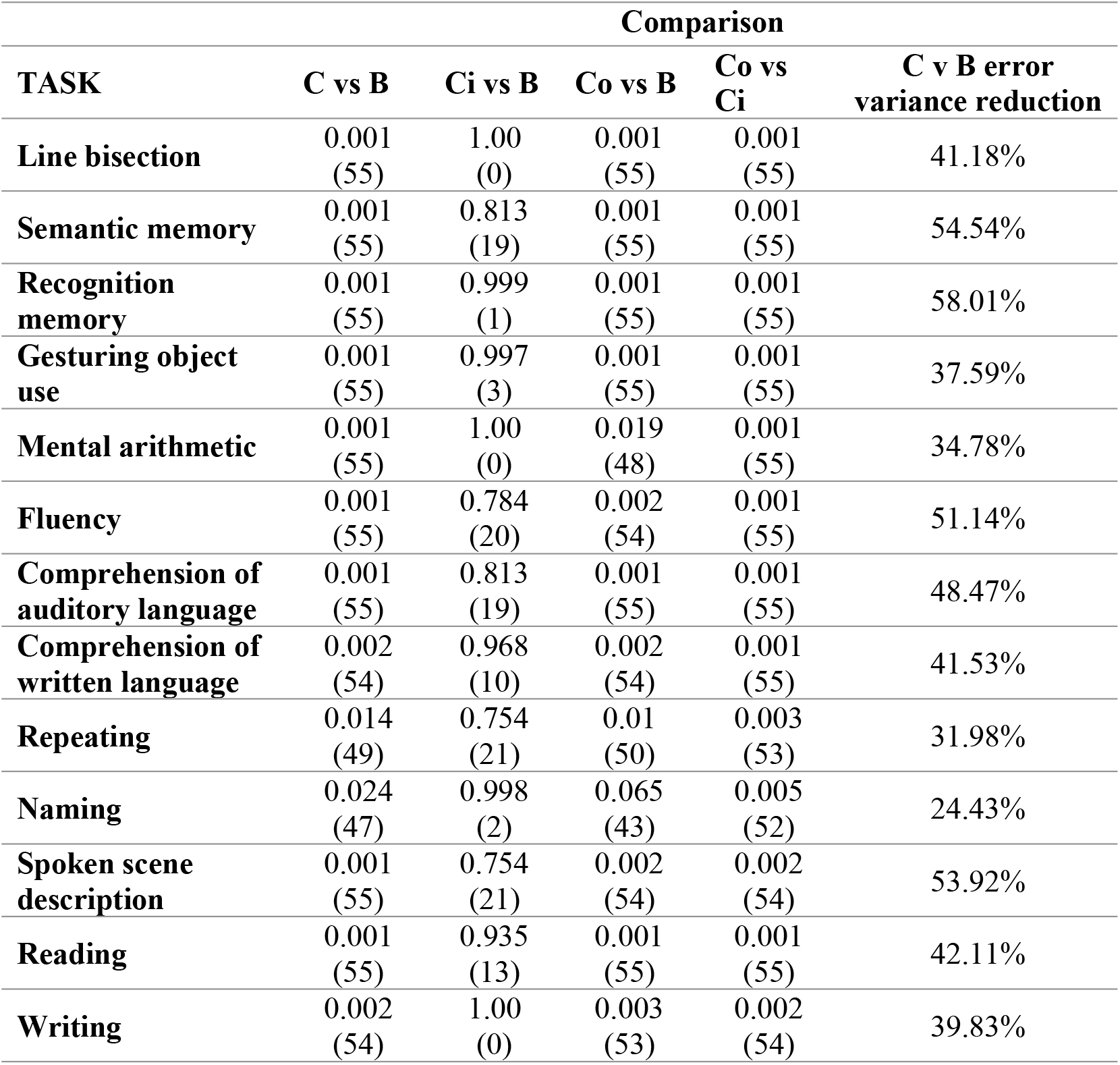
Comparing the predictive performances (errors) of support vector machines driven by different lesion image encodings: B = Binary; C = Continuous; Ci = Continuous-in; Co = Continuous-out. Results are derived from Wilcoxon signed ranks tests, reporting both the p-value and the signed-rank statistic in each case. The final column reports the mean reduction in prediction error distribution variance when using continuous (C) instead of binary (B) lesion images.

**Table 2:** Comparing the predictive performances (errors) of random forests driven by different lesion image encodings: B = Binary; C = Continuous; Ci = Continuous-in; Co = Continuous-out. Results are derived from Wilcoxon signed ranks tests, reporting both the p-value and the signed-rank statistic in each case. The final column reports the mean reduction in prediction error distribution variance when using continuous (C) instead of binary (B) lesion images.

| TASK | Comparison |  |  |  | C v B error<br>variance reduction |
| --- | --- | --- | --- | --- | --- |
|  | C vs B | Ci vs B | Co vs B | Co vs<br>Ci |  |
| **Line bisection** | 0.002<br>(54) | 0.688<br>(23) | 0.001<br>(55) | 0.005<br>(52) | 2.21% |
| **Semantic memory** | 0.116<br>(40) | 0.461<br>(29) | 0.116<br>(40) | 0.097<br>(41) | 1.28% |
| **Recognition**<br>**memory** | 0.003<br>(53) | 0.024<br>(47) | 0.003<br>(53) | 0.032<br>(46) | 3.44% |
| **Gesturing object**<br>**use** | 0.001<br>(55) | 0.947<br>(12) | 0.001<br>(55) | 0.001<br>(55) | 1.85% |
| **Mental arithmetic** | 0.001<br>(55) | 0.539<br>(27) | 0.001<br>(55) | 0.001<br>(55) | 5.24% |
| **Fluency** | 0.002<br>(54) | 0.935<br>(13) | 0.001<br>(55) | 0.001<br>(55) | 3.41% |
| **Comprehension of**<br>**auditory language** | 0.001<br>(55) | 0.423<br>(30) | 0.001<br>(55) | 0.001<br>(55) | 3.24% |
| **Comprehension of**<br>**written language** | 0.007<br>(51) | 0.461<br>(29) | 0.007<br>(51) | 0.014<br>(49) | 2.46% |
| **Repeating** | 0.08<br>(42) | 0.348<br>(32) | 0.065<br>(43) | 0.116<br>(40) | 1.45% |
| **Naming** | 0.007<br>(51) | 0.461<br>(29) | 0.003<br>(53) | 0.001<br>(55) | 4.13% |
| **Spoken scene**<br>**description** | 0.005<br>(52) | 0.993<br>(5) | 0.01<br>(50) | 0.007<br>(51) | 3.64% |
| **Reading** | 0.002<br>(54) | 0.813<br>(19) | 0.002<br>(54) | 0.001<br>(55) | 2.19% |
| **Writing** | 0.019<br>(48) | 0.903<br>(15) | 0.002<br>(54) | 0.005<br>(52) | 1.54% |

Whenever we observed a continuous-binary advantage, we almost always also observed an advantage for continuous-out images as compared to: (a) binary images; and (b) continuous-in images. By contrast, there was almost never an advantage for models driven by continuous-in images; a single exception was observed when predicting recognition memory scores with random forest models. This is evidence that the continuous-binary advantage accrues principally from the availability of graded lesion evidence beyond the binary lesion boundary.

Notably, although the continuous-binary advantage was consistently significant regardless of the machine learning model employed, it was also smaller for random forest models than for support vector machines (median = 2.5% vs. 41.5% reduction in average prediction error distribution variance across all tasks).

## Discussion

Using well-founded model comparison methods, our results suggest that: (a) the use of continuous lesion images confers a predictive advantage when predicting a wide range of language and cognitive outcomes after stroke; and (b) this advantage principally accrues from access to continuous lesion evidence information beyond what might otherwise be considered the binary lesion boundary.

The benefit of continuous lesion information beyond the binary lesion boundary is consistent with growing evidence that the structural integrity of non-lesioned areas contributes to cognitive abilities and recovery after stroke [27, 29, 31-33]. This information may capture a variety of different features, including: (a) subtle damage that was present during the acute phase post-onset; (b) accelerated atrophy due to cerebral diaschisis; and (c) structural changes that enable and sustain post-stroke recovery [27, 28, 34, 35].

One caveat to these results is that, as in all analyses founded on model comparison, there is always a risk that better baseline models might emerge, which extinguish the differences that we have observed here. The random forest model made better predictions than the support vector machine using both binary and continuous lesion images, but the difference between the two models was bigger when both were using binary images. As a consequence, the continuous-binary advantage was therefore smaller for the random forest models than for the support vector machines. The point here is that models that perform even better than the random forest models might reduce the continuous-binary advantage further, by making even better use of the binary lesion images.

Similarly, manual lesion segmentation remains the gold standard in the field, and there remains a risk that models driven by manually segmented lesion images would out-perform those driven by our (algorithmically segmented) binary lesions. We judge this risk to be small because our algorithmic procedure includes humans in the loop, who check that the images correspond to what can be seen by eye. Moreover, the manual segmentation procedure is far from risk free, because its dependence on human raters injects unwanted subjectivity into the results.

Although it may be useful to accommodate continuous lesion information beyond the lesion boundary within a manual lesion segmentation procedure, it is not clear to us how this would be achieved. A solution might be integrate automated lesion identification with manual segmentations of the same lesion that have been approved by multiple raters. We didn’t pursue this route because it would be too resource-intensive and impractical for the scale of our current sample. That said, our algorithmic approach carries resource costs of its own, because it depends on access to scanner-specific images from neurologically normal controls. In brief, we are not claiming that our algorithmic approach is error-free [36] but, without further evidence, we currently believe that its errors should not be systematic.

Despite their caveats, the results presented here do support our previously anecdotal impression, that predictions driven by continuous lesion images out-perform those driven by binary lesions. This advantage appears to generalise across a wide variety of language outcomes scores, and it also applies when predicting non-linguistic cognitive skills after stroke. These advantages add to the confidence with which we can use and interpret the outputs of the ALI toolbox, because they suggest that prognostic accuracy is maximised by considering not just the extremes of the continuous lesion evidence that it supplies, but the whole range of that evidence.

## Ethics

All participants provided informed consent to participate. The study was approved by the Queen Square Ethics Committee.

## Data and code availability

Patient data are available via the conclusion of the data sharing agreement with University College London. Interested researchers should contact Prof. Cathy Price in the first instance. The Automatic Lesion Identification toolbox is an extension for the Statistical Parametric Mapping software, which can be accessed at: SPM Extensions (ucl.ac.uk)

## Author contributions

TMHH and CJP conceived the project, and TMHH implemented it with support from DN, and wrote up the results. MS designed the lesion segmentation algorithm. All authors helped to revise the manuscript.

## Acknowledgements

LORAS team members contributed to the acquisition and analysis of behavioural data. They include: Storm Anderson, Rachel Bruce, Megan Docksey, Kate Ledingham, Louise Lim, Sophie Roberts, and Hayley Woodgate. We are indebted to the patients and their carers for their generous assistance with our research.

